# LANDSCAPE GENOMICS AND DEMOGRAPHY OF CALIFORNIA *AZOLLA*

**DOI:** 10.1101/2025.05.02.651959

**Authors:** Michael J. Song, Carrie M. Tribble, Ixchel González Ramírez, Forrest Freund, Fay-Wei Li, Carl J. Rothfels

## Abstract

*Azolla* was collected and sequenced across California as a part of the California Conservation Genomics Project. We identify three major populations of *Azolla* in the state. Out of these groups, we are able to confidently identify one as *Azolla filiculoides*. The other two taxa are seemingly newly reported to California and are not the same as the taxa currently treated by the Jepson Manual; specifically, they are not members of the *A. microphylla/mexicana* clade. We infer patterns of *Azolla* genetic diversity across the state and the demographic histories of these populations. We find evidence that both isolation by distance and isolation by environment contribute to spatial genetic variation. Both of the two newly reported populations have demographic histories consistent with recent invasions of the state. However, we also discuss alternative possibilities and propose a roadmap for resolving the taxonomy of California *Azolla*.

---

*Azolla* Lam. (Salviniaceae: Salviniales) is one of the most economically important and well-studied genera of ferns, and one of the few fern groups for which there are significant genomic resources available (Li et al., 2018; Song et al., 2025b). It is also one of the most difficult groups for species-level identification, with corresponding challenges with its taxonomy (Metzgar et al., 2007; Evrard & Van Hove, 2004; Madeira et al., 2013), which are further compounded by nomenclatural confusion (Evrard & Van Hove, 2004). The identification of *Azolla* species relies on either microscopic morphological traits or reproductive structures that are difficult to find because the plants often reproduce vegetatively. Even if one can find the putative diagnostic characters, it’s not clear that the corresponding species are monophyletic (Song et al., 2025a) or even if those traits correspond to true species-level units (Evrard & Van Hove, 2004). These challenges exist even in California—despite a long history of detailed characterizations of the California Floristic Province—where up to four different non-invasive species have been reported (*A. filiculoides, A. microphylla, A. mexicana, A. caroliniana*; CCH2 Portal, 2025; Song et al., 2025a), as well the potentially invasive *A. pinnata* (Song et al., 2023), and an additional undescribed species (Li and Rothfels, personal communication).

Currently, the Jepson Manual only recognizes *A. filiculoides* Lam. and *A. microphylla* Kaulf. in California (Jepson Flora Project, 2025). There are only two traits that distinguish these two taxa, both microscopic: *A. filiculoides* has a wider band of cells on the leaf margin than *A. microphylla* and the glochidia are septate in *A. microphylla*, but aseptate in *A. microphylla* (Smith & Murdock, 2012). However, the number of septa has been found to be an unreliable trait, exhibiting variation within species as well as individuals (Lumpkin, 1993). Interestingly, recent studies have identified taxa that are potentially *A. caroliniana* in California (Song et al., 2025a&b). This species is the third species recognized in North America by Flora of North America, alongside *A. filiculoides* and *A. mexicana* (Lumpkin, 1993). *Azolla mexicana* C. Presl is typically treated as a synonym of *A. microphylla* (Smith & Murdock, 2012) and molecular phylogenetic work has supported treating these as a single species (Mezgar et al., 2007).

However, the distinctiveness of *A. caroliniana* from *A. mexicana* has remained controversial. Some studies have supported that the group is one species based on morphological considerations (Edvard and Van Hove, 2004) and RFLP, isozyme, and breeding work (Zimmerman et al., 1989)—asserting that the whole group sister to *A. filiculoides* is a single species that should be named *A. cristata* Kaulf. by priority (for a more detailed description of nomenclatural considerations see Edvard and Van Hove, 2004 and Ahad et al., 2012). On the other hand, Mezgar et al. (2007) concluded that A. *caroliniana* and A. *mexicana/microphylla* are two distinct lineages based on molecular data. There are very few traits that can be used to distinguish the two taxa and only two are given by the Flora of North America: pitted versus non-pitted megaspores and the density of the filosum (Lumpkin, 1993). Complicating matters is that 80 percent of *Azolla* specimens lack these features since they do not contain sporocarps (Lumpkin, 1993). Therefore, despite a great deal of historical sampling, effectively nothing is known about California *Azolla* and the taxonomy of *Azolla* remains an intractable problem.

However, it is possible to study California *Azolla* in a taxonomy-agnostic way by broad molecular sampling with genome resequencing. To this end, we set out to capture the genetic diversity of *Azolla* across the state as a part of the California Conservation Genomics Project (CCGP), a state-funded initiative whose goal is to create genomic resources to guide conservation efforts in the face of climate change (Shaffer et al., 2022; Toffelmier et al., 2022; Fiedler et al. 2022; Beninde et al. 2022). With our broad sampling, we attempted to identify how many distinct populations of *Azolla* there are in the state, identify hotspots of genetic diversity, and infer demographic history of these populations to potentially identify phenomena like recent invasions (Song et al., 2025a). We demonstrate that the information gleaned from landscape genomics studies can be used to generate hypotheses and offer a path to potentially resolving difficult taxonomic problems in cryptic taxa.

## Methods

### Sampling

One hundred and twelve samples of *Azolla* were collected across the state of California. Collection and sequencing protocols and metadata have previously been published (Song et al., 2025a). These samples span the range of *Azolla* in the state (Figure 1 A). We also include in our analyses the published reference genome of *Azolla caroliniana* (Song et al., 2025b). All data generated by CCGP can be found in the NCBI SRA: PRJNA720569.

**FIG. 1.**
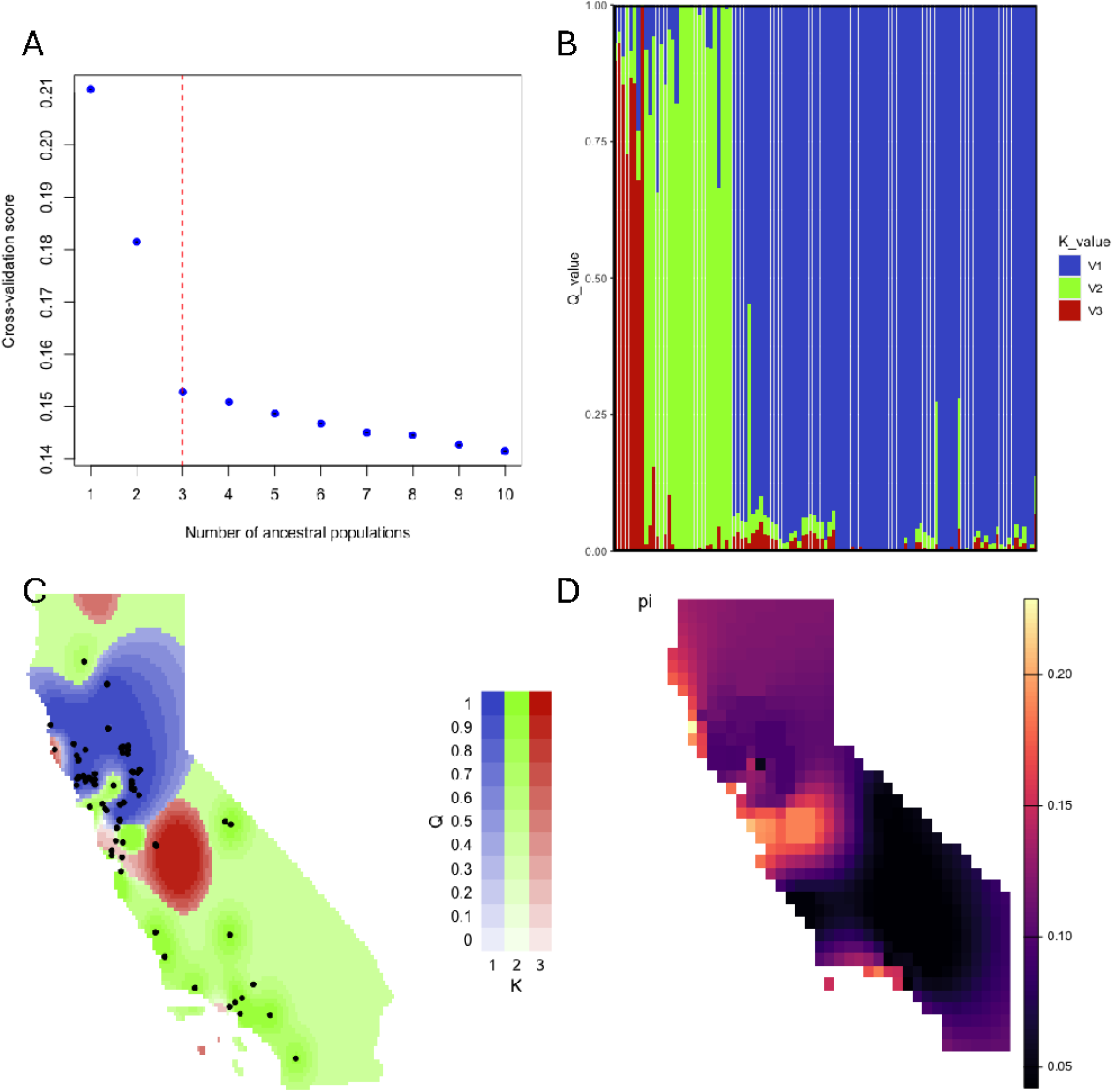
A. Output of the TESS K-testing procedure, which identified that the best k-value obtained is 3. B. Bar plot of ancestry-coefficient values (Q-values) using K=3. Each bar represents one sample and the proportion of the color per sample corresponds to the proportion of ancestry assigned to each cluster. C. Q-values and samples mapped onto California, plotting the maximum Q-value per cell. D. Results of wingen analysis: the genetic diversity statistic *π* mapped onto California.

### SNP Calling

Adapters were trimmed using Trimmomatic (v0.39, Bolger et al., 2014) using the following parameters: LEADING:3 TRAILING:3 MINLEN:36. Samples were then mapped to the *Azolla caroliniana* genome (primary assembly, Song et al., 2025b) using BWA-MEM (0.7.17, Li, 2013). bcftools (v1.9, Danecek et al., 2021) was used to call variants. For the *Azolla* samples, variants were filtered to remove indels and have a QUAL>=30, read depth >10, and the following bias filtering parameters: MQBZ < -3, RPBZ < -3, RPBZ > 3, SCBZ > 3. Out of a total of 44,459,381 SNPs called, ten percent were randomly selected and retained for downstream demography analysis, resulting in a dataset of 861,534 SNPs post-filtering. Population structure analysis was performed using a randomly selected tenth of the demography analysis dataset, retaining a total of 86,134 SNPs.

### Population Structure Analysis

A Landscape Genomic Analysis Toolkit in R (algatr, Chambers et al., 2023) was used to analyze population structure using the TESS method (Caye et al. 2016) and to analyze genetic diversity using moving windows of genetic diversity (wingen, Bishop et al. 2023), both using the default parameters. algatr was used to analyze isolation by distance and isolation by environment using multiple matrix regression with randomization (MMRR, Wang 2013) and generalized dissimilarity modeling (GDM, Ferrier et al. 2007), using the default parameters. alagatr was also used to analyze genotype-environment associations (GEA) using redundancy analysis (RDA, Capblancq & Forester 2021). For all these analyses, we used the top three PCs from a raster PCA on WorldClim database bioclimatic variables. These analyses and visualizations were implemented in R (R Core Team, 2000).

### Inference of Demographic History

We constructed the site frequency spectrum (SFS) for the three populations identified in Song et al. (2025a) using easySFS (Gutenkunst et al., 2009) with the following parameters: -- proj 150,40,18. Population demographic history was inferred from the folded SFS using Stairway Plot v2 (Liu & Fu, 2020) using the default parameters, an assumed mutation rate per site per generation of 1.2e-8, and an assumed generation time of 1 year per generation.

Population size history of *Azolla caroliniana* was also inferred from the reference genome using the Pairwise Sequentially Markovian Coalescent (PSMC) model (https://github.com/lh3/psmc) using the default parameters.

## Results

### Population Structure

We identify three populations of *Azolla* in California (Figure 1 A and B). The larger clade (blue in Figure 1) predominates in Northern California, whereas the other two clades (red and green in Figure 1) are found throughout the state (Figure 1 C). Assessing the distribution of nucleotide diversity (π) across the state, we find that the Bay Area and the coast are hot spots of genetic diversity, while the Central Valley has low values of π, both in the Sacramento Valley and even more so in the San Joaquin Valley. We performed a multiple matrix regression with randomization and found that both geographic distance and one of the PCs of environmental distance were significant—demonstrating that both isolation by distance and isolation by environment contributes to patterns of spatial genetic variation (Table 1). Our generalized dissimilarity modeling results show that geographic distance and all of the PCs are significantly associated with genetic dissimilarity, corroborating the MMRR results (Table 2). In both analyses PC3 was the most significant. The top loadings of PC3 are annual precipitation (0.347), precipitation of coldest quarter (0.325), max temperature of warmest month (0.325), precipitation of wettest quarter (0.323), and precipitation of wettest month (0.322); PC1 and PC2 are loosely more associated with temperature variables. We performed redundancy analysis in order to identify genotype-environment associations and loci that are associated with the three PCs. We found that the top ten most significant SNPs were all associated with PC3. However, a blastn search did not return any hits for any but the most significant SNP on chromosome 22: glyceraldehyde-3-phosphate dehydrogenase B (GAPB) involved in carbohydrate biosynthesis and the Calvin cycle.

**TABLE 1.** Multiple matrix regression with randomization output statistics.

| var | estimate | p | 95%<br>Lower | 95%<br>Upper |
| --- | --- | --- | --- | --- |
| CA_rPCA3 | 0.13 | 0.05 | 0.11 | 0.15 |
| geodist | 0.45 | 0.01 | 0.43 | 0.47 |
| Intercept | 0.00 | 0.89 | -0.02 | 0.02 |

**TABLE 2.** Generalized dissimilarity modeling output statistics.

| predictor | coefficient |
| --- | --- |
| Geographic | 0.67 |
| CA_rPCA1 | 0.66 |
| CA_rPCA2 | 0.21 |
| CA_rPCA3 | 0.80 |

' The percentage of null deviance explained by the fitted GDM model.

### Demography

We inferred population demographic history for three groups of California *Azolla* using SFS and stairway plots. The population in green in Figure 1 (which we call *Azolla filiculoides* in this paper, as per Song et al., 2025a), was estimated to have an effective population size (Ne) of 80,000 and to have experienced population growth over the last thousand generations (Figure 2 A). The population in red in Figure 1 (which we call the “small *Azolla caroliniana* clade” in this paper, as per Song et al., 2025a), which is the smallest clade sampled, was estimated to have an Ne of 100,000 and to have experienced a population contraction around 800 generations ago and a recent population expansion (Figure 2 B). The largest clade sampled, blue in Figure 1 (which we call the “large *Azolla caroliniana* clade” in this paper, as per Song et al., 2025a), was found to have a low Ne of 2,000 and to have experienced very little demographic changes over the last thousand generations, but a recent increase in population size (Figure 2 C).

**FIG. 2.**
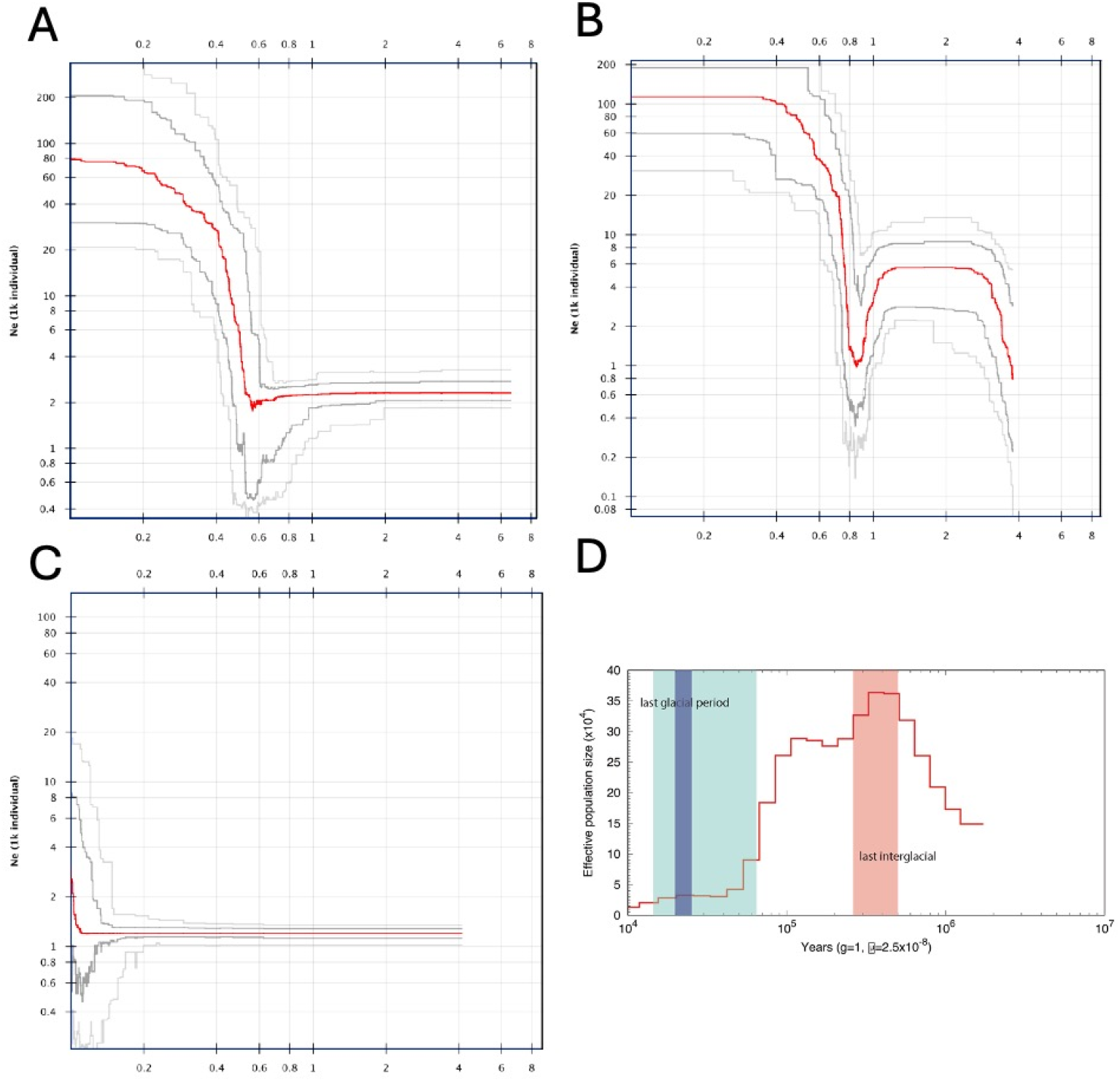
Stairway plots of A. *Azolla filiculoides* B. the small clade of *A. caroliniana* C. the large clade of *A. caroliniana* inferred from the SFS. The x-axis is in units of thousands of individuals; y-axis is in units of thousands of years. D. Stairway plot of the large clade of *A. caroliniana* from the PSMC analysis.

Finally, we estimated population demographic history over a longer time scale using the PSMC model and the reference genome, which was found to be nested within the big *A. caroliniana* clade (Song et al., 2025a). We estimate that over the last 100,000 years, this clade experienced population growth during the last interglacial period and population decline during the last glacial period (Figure 2 D).

## Discussion

Our results, in conjunction with recent publications (Song et al., 2025a&b), indicate that there are at least three distinct *Azolla* taxa in California. Interestingly, we previously did not find a strong association with genetic distance and geographic distance when analyzing the *Azolla* plastome phylogeny inferred from these CCGP data (Song et al., 2025a), but in this study we find a significant contribution of both isolation by distance and isolation by environment. This could be due to the increased amount of information looking at variation across the entire genome rather than just the plastome.

We previously identified that one of the three taxa as most likely *A. filiculoides*, and suggested that the other two clades are potentially *A. caroliniana*, but question whether *A. caroliniana* is monophyletic (Song et al., 2025a&b). This study reveals that these two putative clades of *A. caroliniana* have dramatically different effective population sizes and demographic histories. The small clade, which is genetically similar to *A. caroliniana* samples from eastern North America (Song et al., 2025a&b), was found to have a much larger effective population size than the large clade and to have experienced a recent bottleneck. The large clade, which was collected mostly in Northern California in the Sacramento Valley, was found to have low genetic diversity and a low Ne. It may be that these *Azolla* reproduce sexually less often than the other two California clades, which would also explain the low Ne. The two California clades with high Ne (the small *A. caroliniana* clade and *A. filiculoides*) have maintained a large population size over recent history, whereas the putatively clonal, large *A. caroliniana* clade has the smallest Ne and a recent increase in population size. This points to a possible invasion. While obligate asexual reproduction would dramatically skew estimates of Ne, our samples are facultatively sexual and we even found sporocarps on some of our samples from the putatively more clonal big *A. caroliniana* clade. How facultative sexual reproduction impacts these methods from an empirical perspective, especially when the relative frequency of sex versus asex is unknown warrants further investigation (Hartfield, 2021).

It was recently discovered that *Azolla microphylla*, one of the currently recognized taxa in California, is conspicuously absent from our state-wide sampling (Song et al., 2025a). This raises three potential hypotheses for the origins of these California taxa. First, in the two-invasion hypothesis, a small clade of East Coast *A. caroliniana* may have invaded California, supported by the observed bottleneck in Figure 2B. Independently, the big clade of *A. caroliniana* invaded and established itself in the Central Valley, rapidly spreading clonally, which is supported by the low Ne (Figure 2C). The origins of this second invasion may be from South America, as our previous study (Song et al., 2025b) demonstrated that the large clade was closely related to two International Rice Research Institute (IRRI) samples of *A. caroliniana* collected in Brazil and Uruguay (IRRI 3017, BioSample: SAMN08375392; IRRI 3004, BioSample: SAMN08375393, respectively). Second, in the single-invasion hypothesis, the small clade may represent the plants historically called *A. mexicana/microphylla* in California with those plants being conspecific with the plants from eastern North America (which have historically been referred to *A. caroliniana*). The larger clade is a recent introduction and may be an undescribed species. Alternatively, the large clade could be the historically treated *A. mexicana/microphylla* and the East Coast clade is a recent invasion. Finally, in the no-invasion hypothesis, three clades could represent three native California taxa and taxonomic uncertainty has obscured what has always been present. The molecular evidence linking the small clade to the East Coast and the large clade to South America makes this hypothesis rather unlikely. For it to be true, *A. caroliniana* would have to be two distinct subspecies, one with a native range across the United States overlapping with another that ranges throughout the Americas, and both would have had to have been historically mistaken for one another and described as either *A. mexicana/microphylla* or *A. filiculoides*.

The hypotheses proposed here provide a roadmap for a taxonomic revision of West Coast *Azolla*, as these questions can now be reasonably addressed by further and more expansive low-coverage sequencing. Future studies could sequence historical samples collected across California to see if specimens historically identified as *A. mexicana/microphylla* fall into one of the clades recently described in this study. Likewise, expanded sequencing should be performed on recent samples collected along the West Coast from British Columbia to Baja in order to see if *A. mexicana/microphylla* is still present on the West Coast, just lacking from our CCGP sampling. Our study demonstrates the potential that population-level resequencing can have to address longstanding taxonomic problems in cryptic groups.

## Acknowledgments

We would like to thank our colleagues at CCGP: H. Bradley Shaffer, Erin Toffelmier, Courtney Miller, Merly Escalona, Ruta Sahasrabudhe, Mohan P. A. Marimuthu, Oanh Nguyen, Noravit Chumchim, Eric Beraut, Samuel Sacco, William Seligmann, and Colin W. Fairbairn. This work was supported by the California Conservation Genomics Project, with funding provided to the University of California by the State of California, State Budget Act of 2019 [UC Award ID RSI□19□690224].

## Notes

### Competing Interest Statement

The authors have declared no competing interest.

